# VAL1 as an assembly platform co-ordinating co-transcriptional repression and chromatin regulation at Arabidopsis *FLC*

**DOI:** 10.1101/2021.07.21.453204

**Authors:** Pawel Mikulski, Philip Wolff, Tiancong Lu, Danling Zhu, Caroline Dean

## Abstract

Polycomb (PcG) silencing is crucial for development across eukaryotes, but how PcG targets are regulated is still incompletely understood. The slow timescale of cold-induced PcG silencing at *Arabidopsis thaliana FLOWERING LOCUS C* (*FLC*) makes it an excellent system to dissect this mechanism. Binding of the DNA binding protein VAL1 to an *FLC* intronic RY motif within the PcG nucleation region is an early step in the silencing process. VAL1 interacts with APOPTOSIS AND SPLICING ASSOCIATED PROTEIN (ASAP) complex and POLYCOMB REPRESSIVE COMPLEX 1 (PRC1). Here, we show that ASAP and PRC1 function as co-repressors that quantitatively regulate *FLC* transcription. Upon the shift to cold PRC1-mediated H2Aub accumulates only at the nucleation region, is transiently maintained after transfer back to warm, but unlike the PRC2-delivered H3K27me3 does not spread across the locus. H2K27me3 thus provides long-term epigenetic silencing, whereas H2Aub is a transient repression signal. Overall, our work highlights how a DNA sequence-specific binding protein can act as an assembly platform co-ordinating the co-transcriptional repression and chromatin regulation necessary for Polycomb silencing.

## Introduction

From elegant genetic and molecular analysis in Drosophila the Polycomb group (PcG) proteins have been shown to play a central role in developmental gene regulation^1^. They function in distinct complexes to maintain epigenetic silencing of genomic targets^2^. The most well-studied complexes are POLYCOMB REPRESSIVE COMPLEX 1 (PRC1), which monoubiquitinates the core of histone H2A (H2Aub), and POLYCOMB REPRESSIVE COMPLEX 2 (PRC2), which methylates histone H3 tails at residue 27 (H3K27me). Initially thought to be sequentially recruited (PRC2, then PRC1) many studies have now demonstrated a much more complex scenario with variant and canonical PRC1 ^3,4^, interdependent recruitment of PRC1 and PRC2 and co-operative interactions mediating association to Polycomb Response Elements (PREs)^5–8^. There is also a lack of clarity about target site specificity as Polycomb complexes have been proposed to directly ‘sample’ transcriptionally permissive chromatin sites, rather than be recruited by developmental regulators^9^. Furthermore, the relationship between transcription and chromatin silencing appears to be able to switch between instructive and responsive^10^. Further dissection of the molecular events underpinning Polycomb silencing is therefore necessary to define the core mechanistic principles.

Arabidopsis *FLOWERING LOCUS C (FLC)* is a PcG target central to the process of vernalization ensuring flowering in favourable conditions^11,12^. *FLC* encodes a floral repressor whose expression is high in winter annual Arabidopsis accessions through the activity of FRIGIDA^13,14^. Short cold exposure in autumn/winter results in a relatively rapid transcriptional repression of *FLC*, in a process involving cold-induced *FLC* antisense transcripts called *COOLAIR*^15^. Over the longer timescale of winter, *FLC* is epigenetically silenced through a PRC2-dependent epigenetic switch^16^. This switch occurs at individual *FLC* alleles with a low probability^17^, leading to progressive silencing over the whole plant. A cold-induced step in this switch is the VIN3/VRN5-dependent nucleation of H3K27me3 at an intragenic site that covers three nucleosomes between the transcription start site and the beginning of intron 1^18–20^. When plants return to warm temperatures, H3K27me3 spreads to cover the entire gene, a feature required for the long-term epigenetic silencing throughout the rest of development^17^. The *FLC* system thus provides the opportunity to dissect the components required for a PRC2 epigenetic switch.

Previously, we had shown that the transcriptional repressor VIVIPAROUS1/ABI3-LIKE (VAL1) was a central factor in the switching mechanism, necessary for H3K27me3 nucleation at *FLC*^21^. VAL1 recognizes conserved RY motifs in *FLC* intron 1 with a one bp mutation in the first RY motif sufficient to completely block the vernalization process. The *val1* mutation attenuated *FLC* cold-induced transcriptional repression, but not the longer-term Polycomb memory. VAL1 was shown to interact with components of the APOPTOSIS AND SPLICING ASSOCIATED PROTEIN (ASAP), HDAC complex and PRC1, with components of ASAP interacting with the PRC2 accessory proteins VIN3 and VRN5. This then provided a connection between the sequence at the nucleation region and the machinery for long-term PcG silencing. However, the molecular role of the VAL1 interactors was not known.

Here, we investigate how VAL1 and its interactors ASAP and PRC1 function in *FLC* regulation. Overall, our results suggest that ASAP and PRC1 co-ordinate different steps in *FLC* regulation determining whether and how plants respond to cold, with the sequence-specific transcription factor VAL1 acting as the common recruitment platform. ASAP and PRC1 link co-transcriptional processing with chromatin regulation with PRC1-deposited H2AUb accumulating at the nucleation region during cold exposure. However, this is only maintained at the nucleation region, unlike the PRC2-delivered H3K27me3 which spreads across the locus for long-term epigenetic silencing. Our work reveals how VAL1 functions as an assembly platform to co-ordinate co-transcriptional repression and chromatin regulation at Arabidopsis *FLC*.

## Results

### Mutations of the ASAP complex affect *FLC* expression in warm but not cold-treated plants

To analyse whether ASAP had a role in VAL1-dependent cold-induced FLC silencing we crossed *asap* mutants into a genotype carrying an active *FRI* allele in order to confer a vernalization requirement. The generated FRI *asap* genotypes were analysed for *FLC* expression over a vernalization time course. Plants were harvested before any cold exposure (non-vernalized; NV), immediately after 2, 4 or 6 weeks of cold treatment (2WT0, 4WT0, 6WT0) and after 6 weeks of cold followed by a warm period for 7 days (6WT7). Interestingly, the major effect of the *asap* mutants in both *FRI* and *fri* genotypes was in warm (NV) conditions (Fig. 1a, b). The *asap* mutants did not influence the rate of *FLC* silencing after different cold and post-cold treatments (Fig. 1a). To understand the relationship of VAL1 and ASAP function we generated double mutant combinations in a Col-0 background and analysed in NV conditions. We observed a strong increase in *FLC* expression in warm-grown double mutants of *val1 asap* (Fig. 1b), consistent with VAL1 interacting with many complexes to effect *FL*C silencing. We also found a stronger release of *FLC* silencing in combinations of *asap* mutants (Fig. 1b). As expected with the general correlation of sense and antisense transcription at *FLC*^22^, *COOLAIR* was also upregulated in single *asap* mutants in warm conditions (Fig. S1a). These data suggest that a major role of ASAP at *FLC* is in quantitative transcriptional regulation that sets the level of *FLC* expression in the warm. Consistently, we found an interaction in yeast two hybrid between VAL1 and SIN3A-ASSOCIATED PROTEIN 18 (SAP18), (Fig. 1c,d, S1b), as similarly found in IP-MS with VAL1::VAL1-HA used as a bait^21^. This result is also matching the functional connection between SAP18 and VAL1 reported recently^23^. Given the physical interaction of ASAP components with the PRC2 accessory proteins VIN3 and VRN5, which mediate the cold-induced H3K27me3 nucleation^21^, the lack of an *asap* mutant phenotype during cold and post-cold steps likely indicates functional redundancy with other processes during the cold phase.

**Fig.1.**
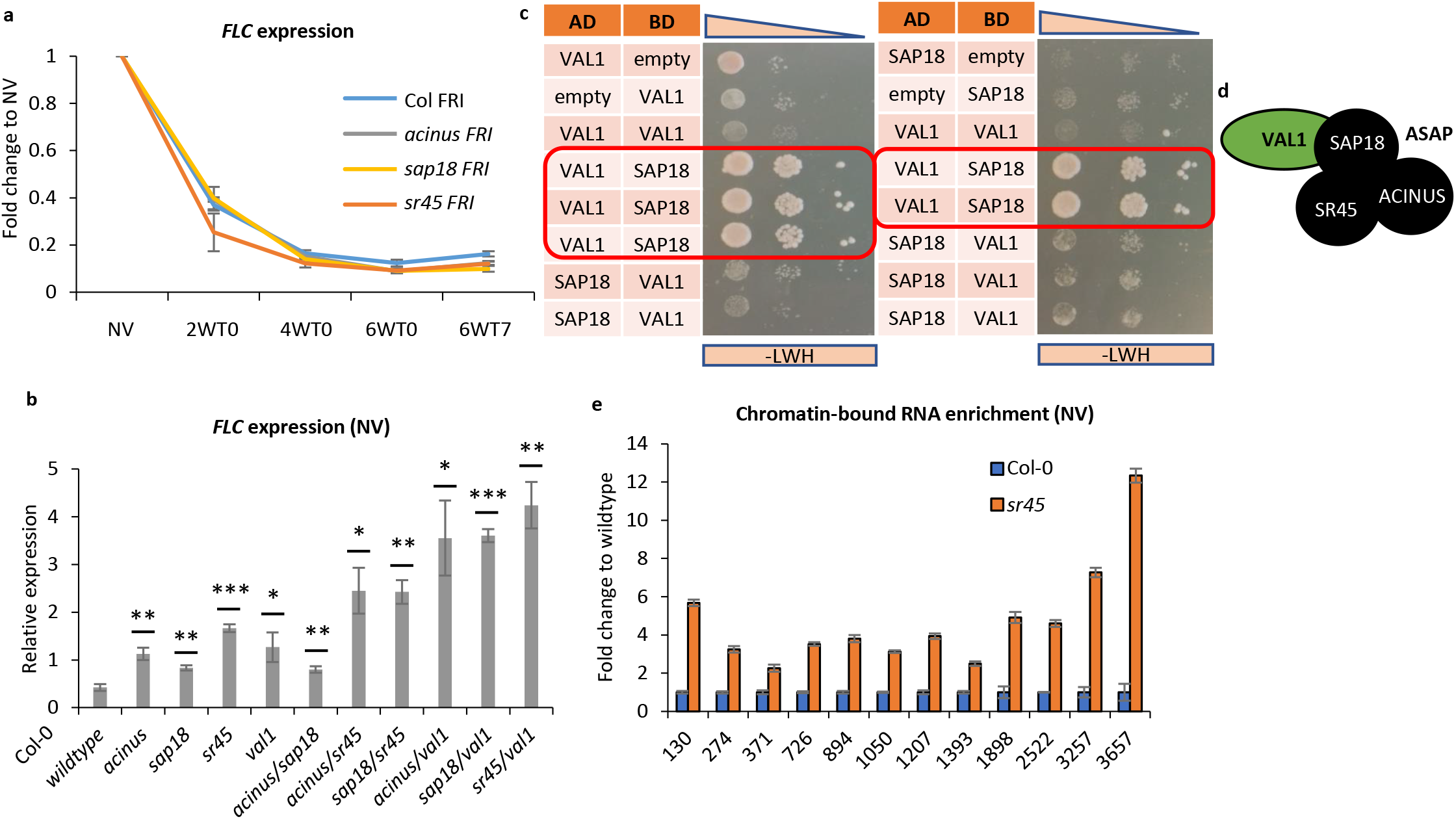
asap mutants affect pre-cold FLC expression only. **a.** FLC downregulation slope during vernalization (fold change to NV). Values were normalized as relative expression to UBC. N = 3 biological replicates, error bars = propagated SEM. **b.** FLC expression in ASAP single mutants and different double mutant combinations within ASAP or with VAL1. Values correspond to mean relative expression to UBC. Statistics are calculated with two-tail Student T-test in comparison to wildtype (* = p≤0.05; ** = p≤0.01; *** = p≤0.001). N = 3 biological replicates; error bars = SEM. **c.** Yeast-two-hybrid of the VAL1-SAP18 interaction. Yeast growth on selective medium SD-LWH is shown. Top panel corresponds to decreasing yeast culture concentration with dilutions: 1/1, 1/5. 1/25 from OD600 = 0.8. AD = pGAD vector backbone with Gal4 activating domain; BD = pGBKT vector backbone with Gal4 binding domain; “empty” = empty vector without insertion as negative control. Red frame depicts protein pairs showing yeast growth over negative controls. **d.** Schematic summary of interaction from a. **e.** Chromatin-bound FLC RNA enrichment over the locus. Shown as a fold change to wildtype. X-axis labels indicate midpoint of the amplicon relative to TSS and its position in FLC locus. N = 3 biological replicates, error bars = propagated SEM.

### SR45-dependent co-transcriptional repression at *FLC*

A role for ASAP outside of Polycomb nucleation at *FLC* led us to analyse a link between co-transcriptional and chromatin regulation, as previously suggested for ASAP at other targets^24^. We selected the *sr45* mutant as an ASAP representative mutant with strong *FLC* upregulation (Fig. 1b) and profiled the co-transcriptional and chromatin changes at *FLC. sr45* increased relative *FLC* transcript levels both in total (Fig, 1b) and chromatin-associated, nascent RNA (Fig. 1e) and *COOLAIR* increased co-ordinately (Fig. S1a). We did not observe changes in *FLC* splicing efficiency at introns 1-3 caused by loss of SR45 or other ASAP components (Fig. S2). However, *sr45* caused higher levels of RNA PolII phosphorylated on serine-2 and serine-5 on its C-terminal domain (CTD) (Fig. 2a), which are associated with active transcription and increased the RNA PolII peak around the *FLC* nucleation region. Furthermore, the balance of active and repressive chromatin modifications was changed in warm grown *sr45* plants, with increased H3K36me3 and decreased H3K27me3 at *FLC* (Fig. 2b), in agreement with the transcriptional upregulation in the mutant. ASAP-dependent co-transcriptional regulation thus correlated with a variety of chromatin features at the *FLC* locus.

**Fig.2.**
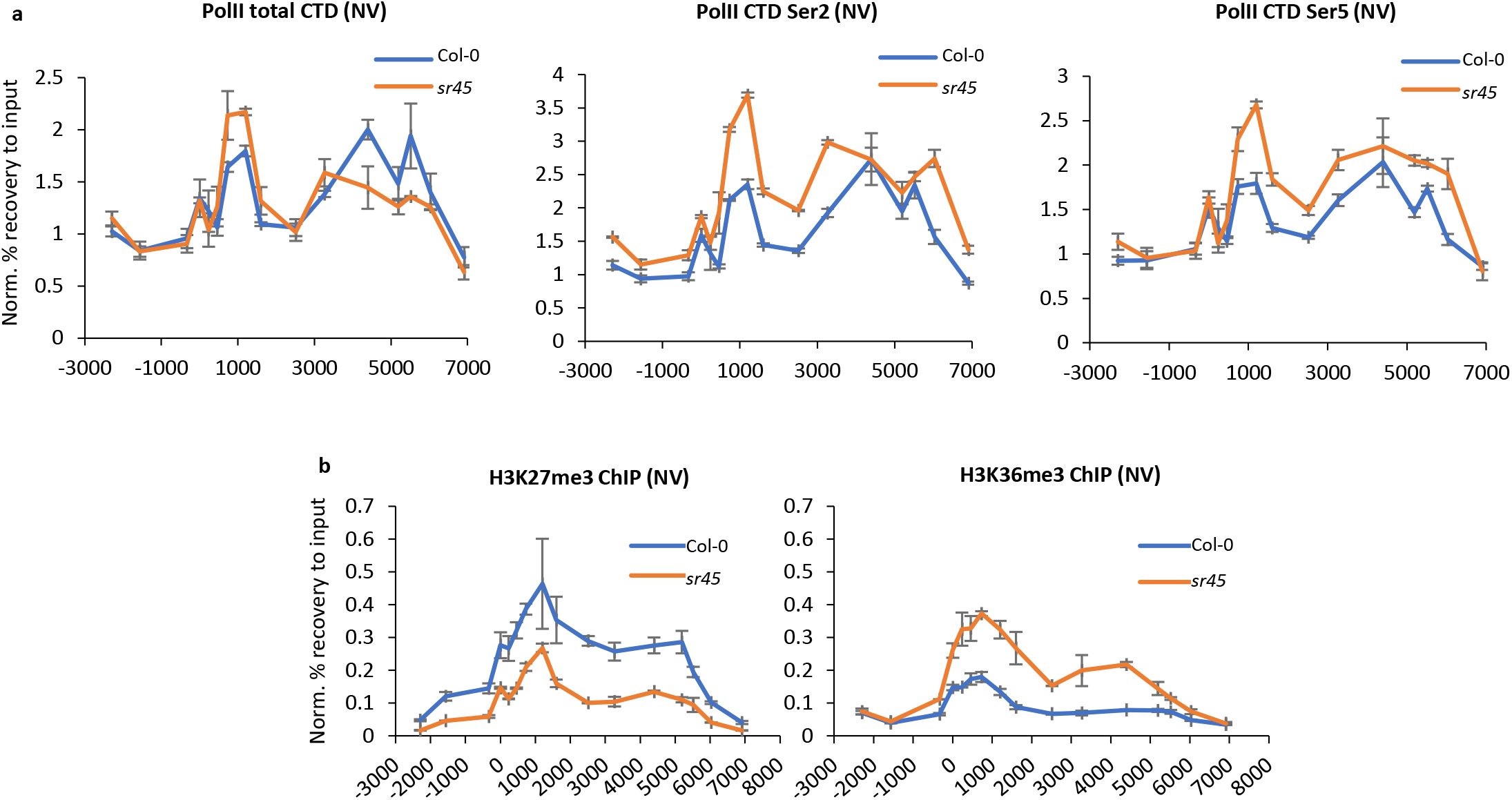
SR45 co-transcriptionally represses *FLC*. **a**. RNA Polymerase II ChIP enrichment. X-axis depicts the midpoint position of amplicon at the *FLC* locus, 0 =TSS. Y-axis corresponds to % recovery to input values normalized to housekeeping gene *ACT7*. NV = non-vernalized; n = 3 biological replicates; error bars = SEM. **b**. SR45-dependent change of active-repressive histone marks. X-axis depicts midpoint position of amplicon at the *FLC* locus. Y-axis corresponds to % recovery to input values normalized to H3 enrichment. NV = non-vernalized; n = 3 biological replicates; error bars = SEM.

### VAL1 affects specific nucleosome dynamics

A role for ASAP on co-transcriptional processes implied that its interactor, VAL1, may also have a similar function, especially given its impact on the active/repressive chromatin modifications^21^. To test this, we analysed chromatin-association of *FLC* RNA in warm-grown *val1* seedlings as a readout for bona fide *FLC* transcription. Like the relative increase in total RNA^21^, and similar to *sr45* (Fig. 1e), *val1* increased chromatin associated *FLC* RNA (Fig. 3a). Given the VAL1 binding site is downstream of the *FLC* promoter we asked if such transcriptional changes might be accompanied by changes in nucleosome dynamics^25^. We adapted the histone salt fractionation protocol that preferentially extracts nucleosomes recently disrupted by either transcription or remodelling^25^ and analysed *FLC*. The fraction of low-salt extractable nucleosomes decreased at the +1 and +2 nucleosomes in cold-treated samples (Fig. S3), agreeing with the previous observation of cold-induced nucleosome stabilization^19^. Furthermore, we observed that *val1* increased the low-salt extractable fraction for nucleosomes +1 and +3, but not for nucleosome +2 (Fig. 3b, c). Nucleosomes +1 and +3 correspond to the edges of the H3K27me3-enriched nucleation region found in *FLC* after cold exposure^26^. These VAL1-dependent changes were confined to chromatin extracted from warm-grown *val1* seedings and were not found in cold-treated samples (Fig. S3). Interestingly, they were not connected to gross changes in chromatin accessibility as judged by FAIRE (formaldehyde-assisted isolation of regulatory elements, a technique using nuclease hypersensitivity to identify open and therefore potentially regulatory chromatin) (Fig. 3d). Overall, these data connect VAL1-induced co-transcriptional quantitative regulation to nucleosome dynamics at the edges of the *FLC* nucleation region, a region where RNA PolII occupancy is high.

**Fig.3.**
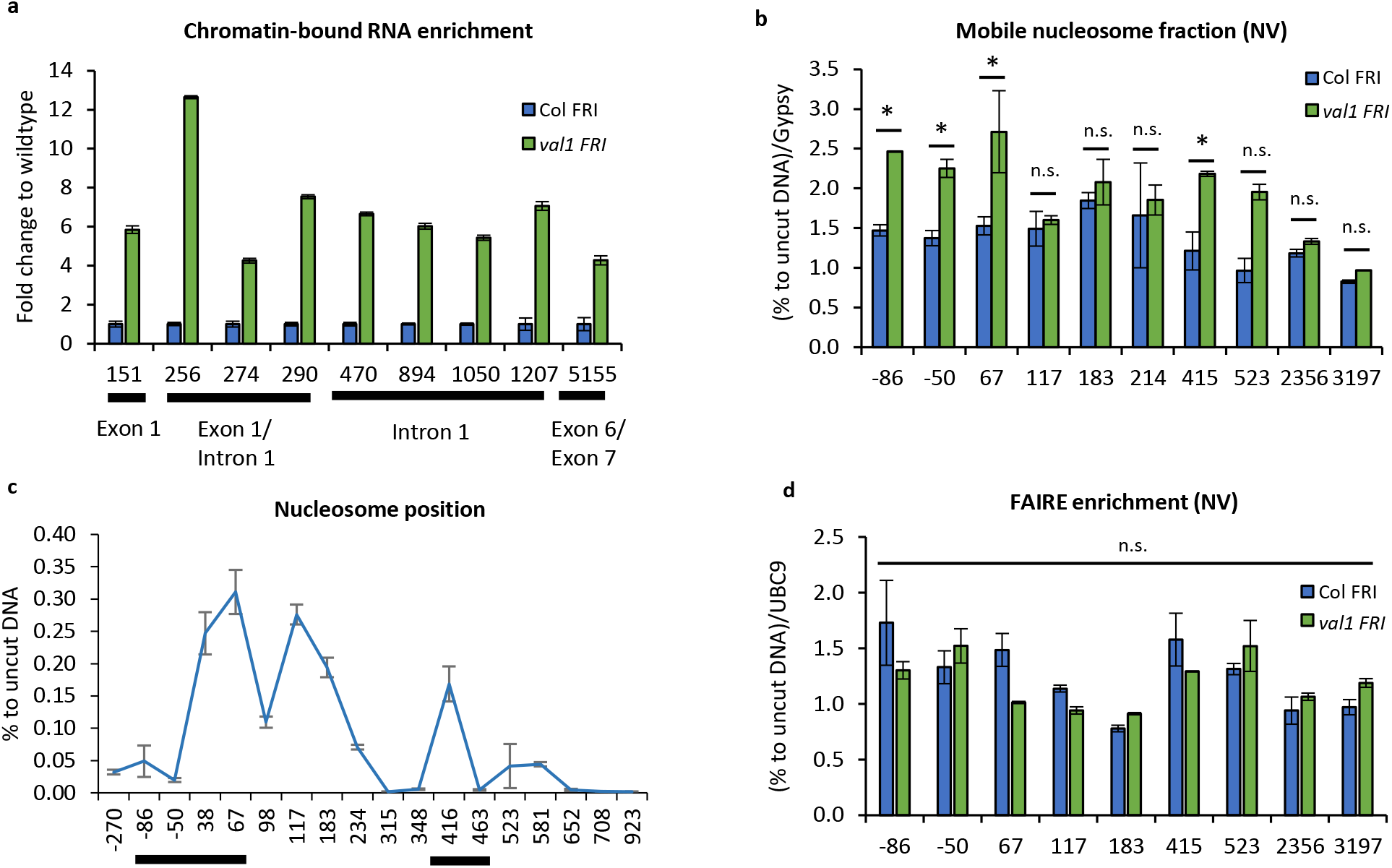
*FLC* chromatin regulation by VAL1. **a**. Chromatin-bound *FLC* RNA enrichment over *FLC* locus. Shown as a fold change to wildtype. X-axis labels indicate midpoint of the amplicon relative to TSS and its position in *FLC* locus. N = 3 biological replicates, error bars = propagated SEM. **b**. Low salt-extractable nucleosome fraction over *FLC*. Shown as % recovery to uncut DNA and fold to AT4G07700 (Gypsy-like transposon). X-axis labels indicate beginning of amplicon relative to TSS, lines below axis’ label indicate regions of increased low-salt nucleosome extractability in *val1* shown in b. N = 3 biological replicates; error bars = SEM; asterisk = p<0.05 from two-tail Student T-test in comparison wildtype vs *val1*; n.s. = non-significant; NV = non-vernalized. **c**. Nucleosome position at *FLC* in wildtype. Results from MNase-qPCR assay shown as % to uncut DNA. X-axis indicate beginning of amplicon relative to TSS. N = 2 biological replicates, error bars = SEM. **d**. FAIRE enrichment at *FLC*. Shown as % recovery to non-crosslinked sample (UN-FAIRE) and fold to UBC9. X-axis labels indicate beginning of amplicon relative to TSS. N = 2 biological replicates; error bars = SEM; NV = non-vernalized.

### H2Aub accumulates during cold at the *FLC* nucleation region but does not spread like H3K27me3 upon return to warm

To begin to investigate the functional significance of the VAL1-PRC1 interaction we undertook analysis of H2AUb accumulation at *FLC* on the vernalization-sensitive genotype ColFRI after different cold exposure. Reduced *FLC* expression correlated with increasing H2Aub levels as plants were exposed to progressive cold (Fig. 4a). H2Aub accumulated at the *FLC* nucleation region with dynamics similar to PRC2-induced H3K27me3 nucleation. To establish whether the H2Aub correlated with *FLC* transcriptional activity we then analysed lines with or without FRIGIDA (FRI), which is the main determinant distinguishing vernalization requirement or rapid-cycling habit. In the rapid-cycling Col-0, H2Aub levels at *FLC* were much higher than in Col-FRI reflecting the ∼ 20x reduction in *FLC* transcription (Fig. 4b). These results suggest that H2Aub is inversely correlated with transcription and PRC1 has a typical transcription repression function at *FLC*, rather than a context-dependent role in transcriptional activation^3,4^.

**Fig.4.**
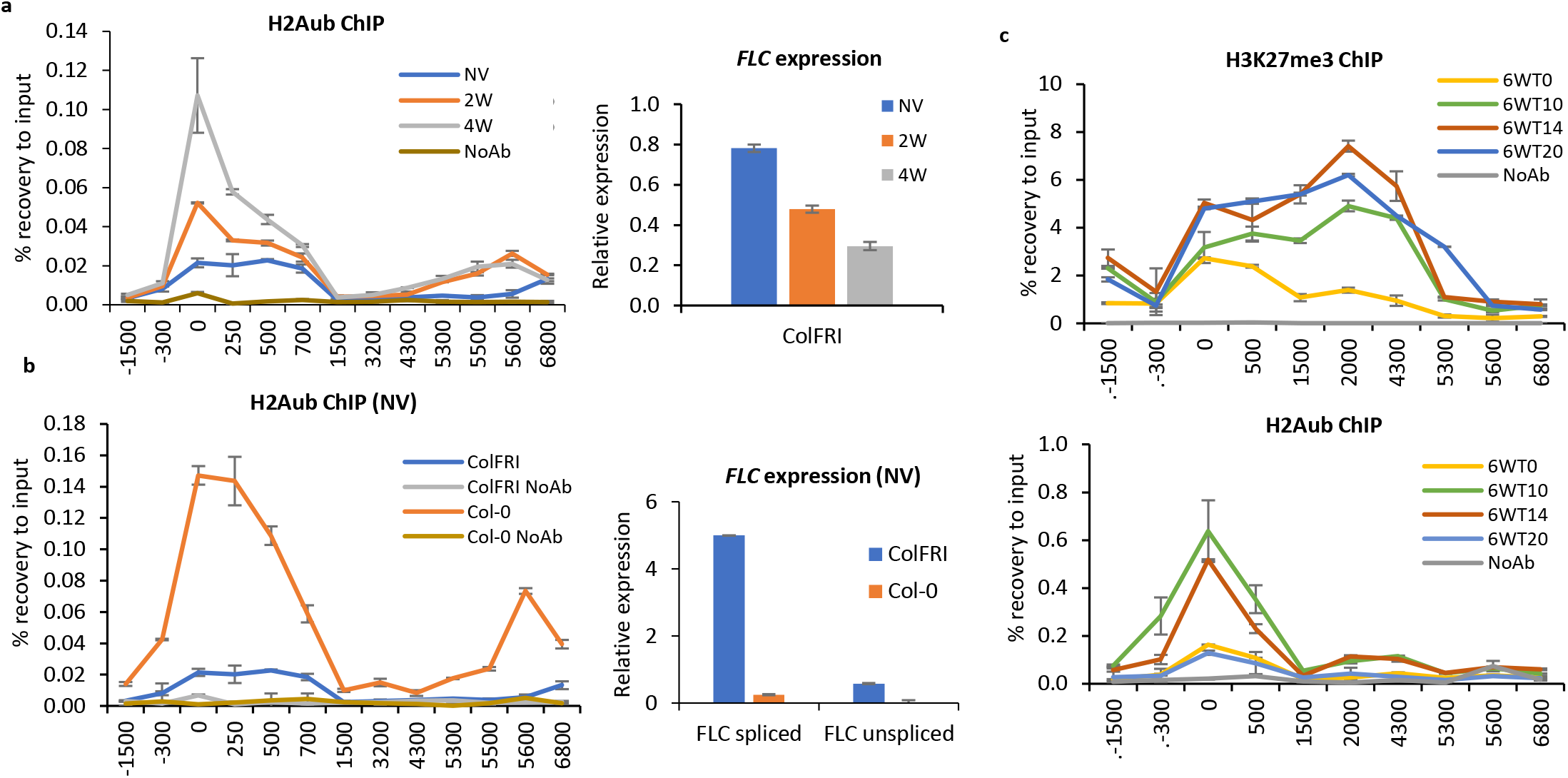
H2Aub dynamics at *FLC*. **a**. H2Aub chromatin -IP showing strong enrichment of H2AUb at the 5’ end of *FLC* in ColFRI correlating with increased transcription in non-vernalized (NV) conditions. X-axis show midpoint amplicon at *FLC*. N = 3 biological replicates; error bars = SEM; NoAb = no antibody negative control. *FLC* expression in wildtype backgrounds in NV is depicted in graph on the right. N = 3 biological replicates, error bars = SEM. **b**. H2Aub chromatin -IP correlating cold-induced decrease in *FLC* expression with decreased H2AUb at the 5’ end of *FLC*. X-axis show midpoint amplicon at *FLC*. Graph on the right shows unspliced *FLC* expression during vernalization timecourse. NV = non-vernalized; Nominal-W = number of weeks of cold; N = 3 biological replicates; error bars = SEM; NoAb = no antibody negative control. **c**. Histone mark behaviour post-cold. H3K27me3 and H2Aub enrichment over *FLC*. X-axis show midpoint amplicon at *FLC*. N = 3 biological replicates; error bars = SEM; 6W = 6 weeks of cold treatment; T-nominal = number of days post-cold; NoAb = no antibody negative control.

We then tested how H2Aub changes in warm conditions post-cold, when *FLC* chromatin transitions into a stably repressed state. We measured H2Aub and H3K27me3 at *FLC* during the vernalization timecourse^26^. As expected, H3K27me3 accumulated in the nucleation region during cold exposure (6W+day 0) and spread across the whole gene body after transfer back to warm (6W+day 10/14/20) (Fig. 4c). H2Aub accumulated in the nucleation region during cold, similar to the H3K27me3 dynamics, and continued to increase for 10 and 14 days after transfer back to the warm but did not spread across the locus (Fig. 4c). Interestingly, 20 days after transfer back to the warm, the H2Aub peak at the nucleation decreased back to levels seen at the end of 6W cold. These dynamics closely match the occupancy of the VRN5 accessory protein at the nucleation region^17^.

### VAL1, PRC1 and NDX physically interact and are required for H2Aub accumulation at the *FLC* nucleation region

Genetic dissection of the role of H2AUb at *FLC* is complex as the multiple Arabidopsis PRC1 components are functionally redundant and single PRC1 mutants do not show a changed *FLC* expression relative to wildtype^27,28^ (Fig. 5a). Loss of all three BMI subunits (A, B, C) de-represses *FLC* expression in warm-grown Col-0 plants up to 25 fold^29^. Higher order PRC1 mutant combinations are seedling lethal or show severe developmental aberrations^27,30,31^. In addition, Arabidopsis PRC1 has a range of partners central to its involvement in different developmental pathways^32^. However, to pursue the consequence of the VAL1-PRC1 interaction we analysed *FLC* expression differences in *val1 bmi1B* line. *val1 bmi1B* did not increase *FLC* expression above *val1* alone in any condition tested (Fig. 5b, c) or further influence *FLC* H2Aub levels in warm-grown plants (Fig. 5d). In addition, *val1 bmi1B* did not show *FLC*/*COOLAIR* reactivation post-cold (Fig. S4). This was also true for the combination *val1 ring1A* (Fig. S5). We did find an additive effect of the *val1 bmi1B* combination on *COOLAIR* expression (Fig. S6), so for this aspect of *FLC* regulation other PRC1 components do not completely cover loss of BMI1B function.

**Fig.5.**
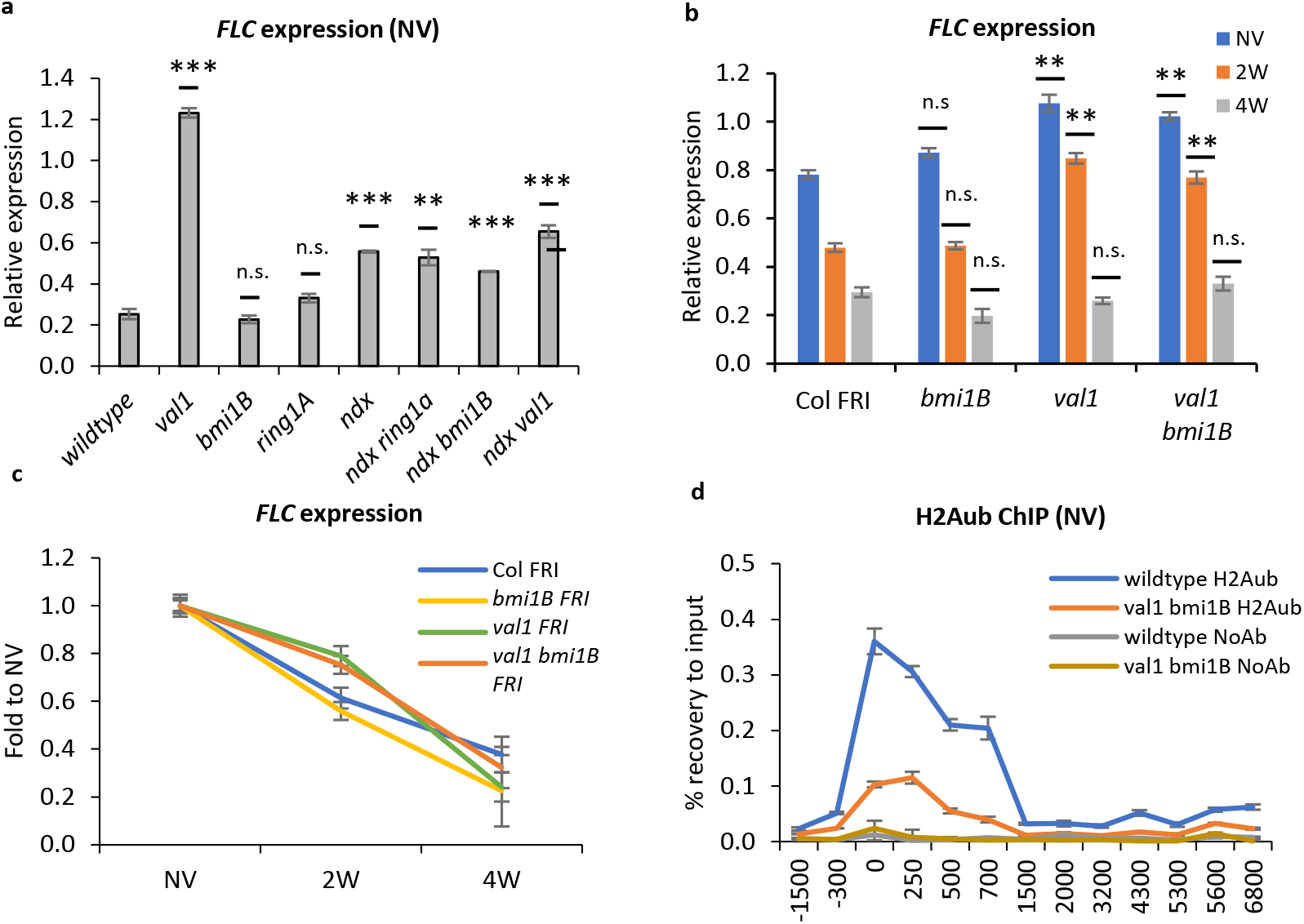
VAL1, BMI1B and NDX effect on *FLC* expression. **a**. *FLC* expression in mutants of VAL1, PRC1 and NDX for *FLC* RNA. Values correspond to mean relative expression to geometric mean of UBC and PP2A. Statistics were calculated with two-tail Student t-test in comparison to wildtype. N = 3 biological replicates; error bars = SEM. **b**. *FLC* expression level. Represented as relative expression to housekeeping genes (see methods). Statistics represent two-tail Student t-test in comparison to the wildtype, N = 3 biological replicates, error bars = SEM, NV = non-vernalized, W = number of weeks of cold. **c**. The same data as in **b**. presented as *FLC* downregulation slope during vernalization (fold change to NV). **d**. H2Aub enrichment at *FLC* in wildtype and *val1 bmi1B*. N = 3 biological replicates; error bars = SEM; NoAb = no antibody negative control.

We undertook further IP-MS experiments with VAL1-HA as a bait and found a robust interaction between VAL1 with other proteins (Table S1), including PRC1 members, as well as NODULIN HOMEOBOX (NDX) (Fig. 6a). Interestingly, NDX has been previously implicated in single-stranded DNA recognition and stabilization of the R-loop at *FLC*^33^. Using yeast-two-hybrid analysis we found that NDX does not interact directly with VAL1 but instead interacts with RING1A and RING1B (PRC1 components) (Fig. 6b, c, S7), as also found recently^34^. Genetic analysis between *val1, prc1* mutants and *ndx1* revealed that NDX is required for the *FLC* upregulation caused by *val1*, and loss of PRC1 does not affect the consequences of *NDX* deficiency (Fig. 5a); *COOLAIR* was upregulated similarly to *FLC* in the mutants (Fig. S8). Furthermore, we also showed that NDX, VAL1 or VAL1-recognized RY motifs are necessary for full H2Aub enrichment at *FLC*, as well as H3K27me3 (Figs. 6d, e, S9 and as published in^21^). NDX has been shown to associate with the 3’ region of *FLC* and stabilize R-loops^33,35^, so it will be important to establish if these genetic interactions point to an additional NDX/R-loop interaction in the *FLC* nucleation region or are the result of the gene loop^36^.

**Fig.6.**
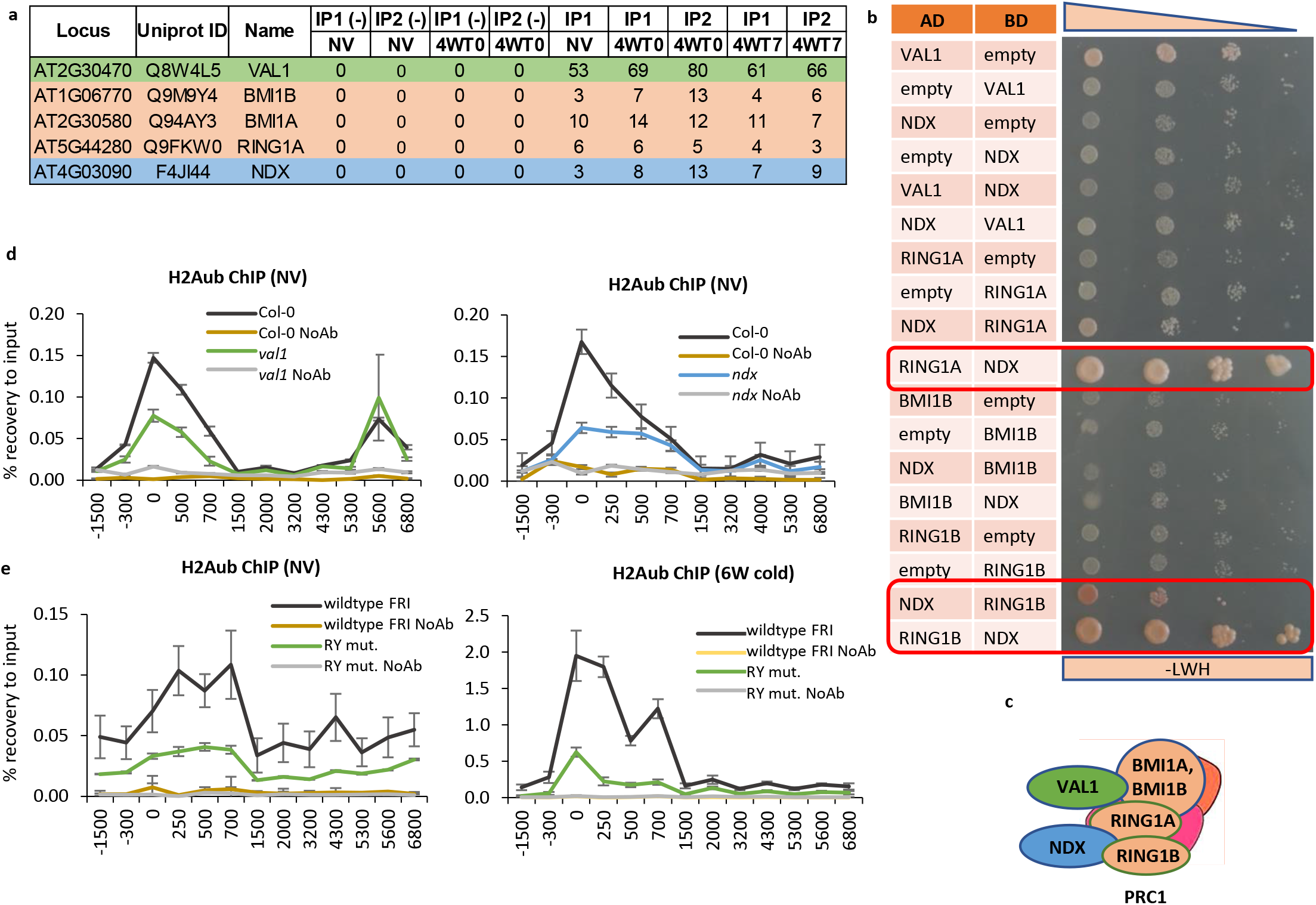
VAL1, PRC1 and NDX physically interact and are required for H2Aub accumulation at the *FLC* nucleation region. **a**. IP-MS/MS results for PRC1 and NDX from crosslinked VAL1-HA co-IP. Numbers corresponds to unique peptides. IP-numeral = biological replicate; NV = non-vernalized; Numeral-W = number of weeks of cold; T-numeral = number of days post-cold; (-) = negative control. **b**. Yeast-2-hybrid results for PRC1-NDX interaction. Yeast growth on selective medium (-LWH) is shown. Top panel corresponds to decreasing yeast culture concentration. AD = pGAD vector backbone with Gal4 activating domain; BD = pGBKT vector backbone with Gal4 binding domain; “empty” = empty vector without insertion as negative control. Red frame depicts protein pairs showing yeast growth over negative controls (pairs with empty vector). precipitated with anti-HA antibody. **c**. Schematic summary of interactions from **a**. and **b. e**. and **f**. H2Aub ChIP results. X-axis represents midpoint of the amplicons over *FLC*. NoAb = no antibody negative control; N = 3 biological replicates for H2Aub IPs and 3 qPCR technical replicates for NoAb controls; error bars = SEM.

H2Aub enrichment at *FLC* was unaffected in mutants of the core PRC2 component VERNALIZATION 2, or the PRC2-accessory protein VERNALIZATION 5 in non-vernalized conditions but was slightly reduced after 6 weeks cold exposure (Fig. S10). The lack of difference in NV conditions is in agreement with H2Aub being generally independent of LHP1 and PRC2 function in Arabidopsis, as previously reported^37^. However, the small effect after cold potentially connects close functioning of VAL1, NDX and PRC1 in *FLC* co-transcriptional repression to PRC2 nucleation and chromatin silencing.

## Discussion

How transcriptional repression links to epigenetic silencing is far from understood. Cell type-specific or developmentally induced transcriptional repressors bind different Polycomb complexes but direct recruitment models are too simplistic^9,10^; linear genetic hierarchies have not been established and functional output of PREs has been shown to be heavily dependent on the chromatin context^38^. Our findings that VAL1 acts as an assembly platform co-ordinating ASAP and PRC1 activities that occur at different stages in the *FLC* regulation helps explain some of this complexity. It also raises interesting parallels to the non-linear recruitment activities of the PcG proteins at Drosophila PREs^7,39,40^, with ASAP co-transcriptional repressors and PRC1 components likely redundantly establishing the chromatin environment required for robust PRC2 silencing (Fig. 7).

**Fig.7.**
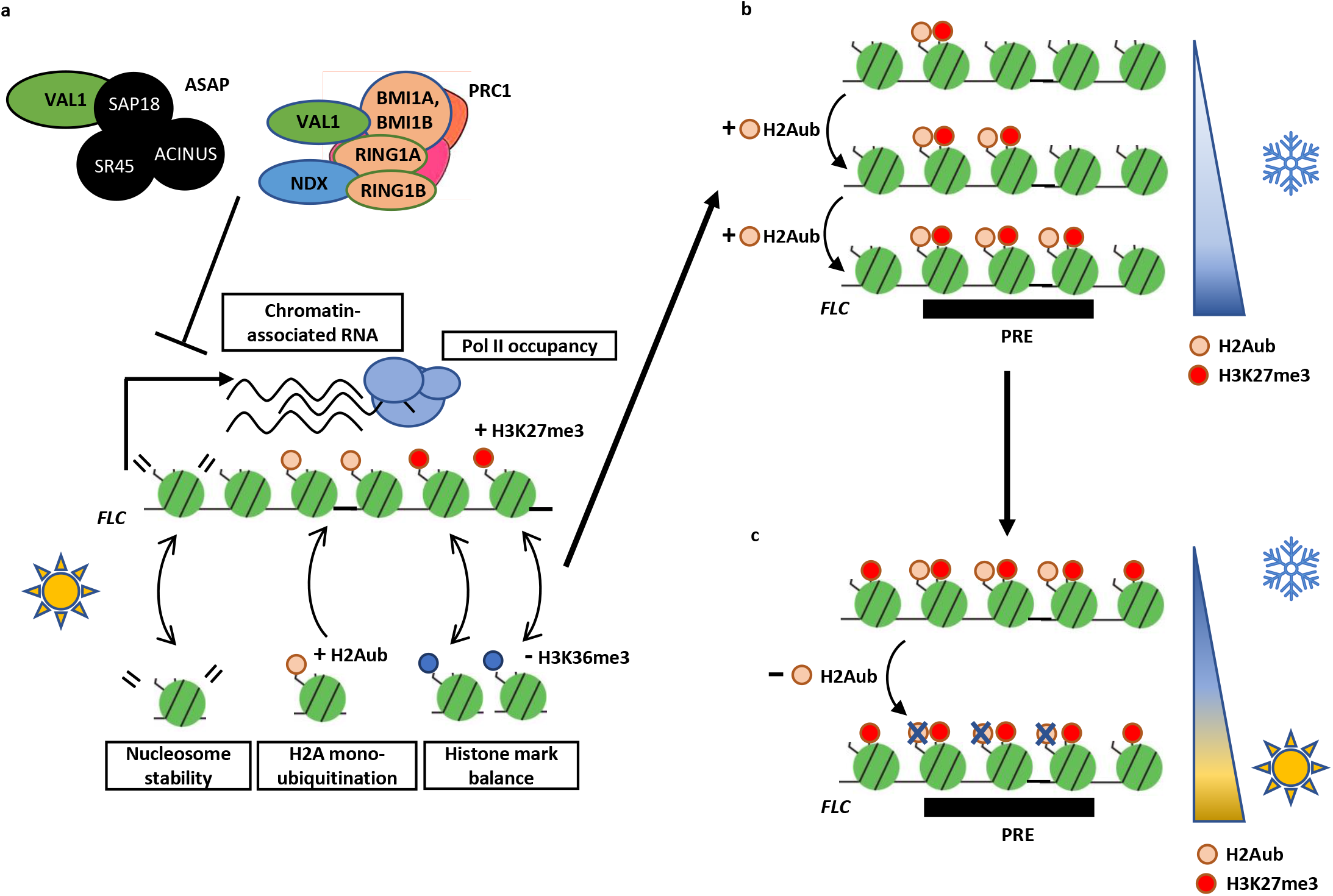
Functional relationship of ASAP, PRC1 and PRC2 in *FLC* regulation. **a**. ASAP and PRC1 act as co-repressors that regulate *FLC* chromatin features and downregulate *FLC* expression in warm. **b**. Upon shift to cold, PRC1-mediated H2Aub accumulates at *FLC* PRE along with H3K27me3. H2Aub and H3K27me3 enrichment gradually increases along with cold duration. In post-cold, H3K27me3 spreads to *FLC* gene body, whereas H2Aub remains localized to the *FLC* PRE. **c**. At the later post-cold timepoints, H3K27me3 is stably maintained over the *FLC* gene body whereas H2Aub is removed from the chromatin. H2K27me3 thus provides long-term epigenetic silencing, whereas H2Aub is a transient repression signal.

ASAP and PRC1 have been documented to have very different roles in gene regulation in different organisms. ASAP has previously been shown to play a role as part of the spliceosome and exon junction complexes in many organisms^24,41^, whereas, PRC1 functionality is associated with H2AUb and transcriptional silencing^42^. In plants, ASAP has been associated with regulation of innate immunity^43^ and abiotic responses^44^ and considered to function through altering RNA metabolism or alternative splicing^45,46^. However, we did not detect any effect of a *sr45* mutation on alterative splicing or splicing efficiency of the *FLC* sense strand introns. Instead, we found changes linked to transcriptional repression – RNA PolII post-translational modifications and classic histone modifications associated with Polycomb target ON or OFF states. We thus propose that SR45, as an ASAP component, influences a co-transcriptional step that may function to influence RNA PolII complex composition and thus the chromatin modifications deposited during transcription. Since there is physical linkage of VIN3 and VRN5^21^ to ASAP components, the ASAP complex is likely to function at *FLC* both during growth in the warm, as judged by mutant phenotypes, and redundantly with other functions in *FLC* silencing during cold exposure. Changed co-transcriptional regulation with RNA PolII both involved in histone modification deposition, but also sensitive to histone modifications on the chromatin template, would seem a probable mechanism underlying the instructive/responsive transcription-chromatin relationship.

The PRC1-mediated H2AUb modification has been associated with transcriptional regulation and epigenetic stability in Drosophila and mammalian cells^42^. At *FLC*, the PRC1-H2AUb is associated with transcriptional repression, either in low *FLC* expressing genotypes (Col-0) in warm conditions, or in response to cold in the high *FLC* expressing Col-FRI genotype. In this respect it appears to function at *FLC* more as a variant PRC1 rather than a canonical PRC1. The timescale of the VAL1-PRC1-mediated H2Aub accumulation at the *FLC* nucleation region during cold exposure suggests an intermediary role between the transcriptional silencing determined by ASAP-VAL1 and the long-term PRC2 epigenetically stable spread state. However, the genetic redundancy makes PRC1 functionality hard to tease apart so we were unable to dissect the interdependency of PRC1-delivered H2AUb and PRC2-delivered H3K27me3 in the epigenetic switch at *FLC*. Therefore, how VAL1 interaction with ASAP and PRC1 is co-ordinated spatially and temporally at the *FLC* nucleation region remains to be fully elaborated.

VAL1 interacts with ASAP via SAP18 and PRC1 but has also been shown to interact with LHP1^32^, histone deacetylases (HDA9, HDA6)^47,48^ and Mediator complex component (MED13)^47^. It will be important in the future to understand how one protein can mediate all these interactions and whether they are reinforced through multivalent interactions with other proteins at the nucleation region, like the Drosophila PRE situation. The original intronic mutation in the RY motif caused relatively little warm mis-regulation of *FLC*^21^, just attenuation of the cold-induced transcriptional silencing without a subsequent effect on stability of the silenced state. The stronger phenotype of the VAL1 interactors in the warm argues that they also function independently of VAL1. The lack of phenotype in the cold suggests that functionally redundant, cold-specific regulators are still to be identified. These are likely to involve the antisense transcripts at *FLC* (*COOLAIR*) that contribute to transcriptional silencing during cold exposure, which associate with the chromatin at the 5’ end of *FLC*, and in which natural variation has a functional consequence^49,50^.

Overall, we interpret our data as showing that VAL1 functions as an assembly platform to co-ordinate co-transcriptional repression and chromatin regulation at Arabidopsis *FLC*. We envisage that transcription is downregulated by multiple repressor complexes (ASAP and PRC1) either linked to RNA PolII and interacting with VAL1 at the nucleation region or assembled at the nucleation region. These appear to function independently in the warm, with H2AUb playing a major role in the cold associated with *FLC* silencing, alongside PRC2 deposition of H3K27me3. Given the parallels with the Drosophila PcG mechanism the activities of these many protein complexes are likely to function cooperatively to influence the different steps in the mechanism. Further elucidation will require detailed analysis of the co-transcriptional silencing, chromatin modification and nucleosome remodelling.

## Material and methods

### Plant material

All mutants and transgenic lines were either in Col-0 or FRI^sf2^ background^21,50^. Unless otherwise stated, experiments in non-vernalized conditions used Col-0 as a wildtype. The analyses in vernalization timecourse used Col FRI^sf2^ (Col FRI) as a wildtype to capture dynamics of cold-induced *FLC* expression and chromatin changes. Mutant alleles were used as described previously: *bmi1B* (*drip1-1*), *bmi1A* (*drip2-1*), *ring1A* (GK-293A04), *ring1B* (SALK_117958), *ndx1-4, flc* (*flc-2*), RY motif (*vrn8*), *val1*-2, *vin3-4, vrn2-1, vrn5-8, sap18* (SALK_02363C), *sr45* (SALK_018273), *acinus* (SALK_078554)^17,21,33,34,51^. Double mutants were generated by crosses between homozygous single mutant lines and selected by PCR-based genotyping using primers listed in Table S2.

### Growth conditions

Seeds were surface sterilized and sown on MS-GLU (MS without glucose) media plates and stratified at 4°C in the dark for 2 days. For non-vernalized (NV) conditions, seedlings were grown for 14 days in long-day conditions (16 h light, 8 h darkness with constant 20°C). For vernalization treatment, seedlings were grown at 5°C in short-day conditions (8 h light, 16 h darkness with constant 5°C) for number of weeks indicated in the respective experiment description.

### Protein co-immunoprecipitation

4 g of Arabidopsis seedlings were crosslinked and ground in fine powder by liquid N2, following ChIP protocol^21^. Immunoprecipitation of protein complexes was done as described in^17^ using Protein A beads (Thermo Fisher Scientific, #10008D) coupled with HA antibody (CST, #3724). IP product was resuspended with 2X NuPAGE sample buffer (Thermo Fisher Scientific, #NP0007) and denatured at 70°C for 15 min with 100 mM DTT. The final samples were run on NuPAGE Bis-Tris gels (Thermo Fisher Scientific, # NP0321) and subjected to in-gel digestion.

Protein bands corresponding to heavy and light chains of antibody were excised from NuPAGE gels. The rest of gels were combined and performed in-gel digestion as described previously^52^. Briefly, gel pieces were destained and then incubated with 10 ng/μl trypsin (Promega, #V5111) overnight at 37°C. Peptides were extracted by incubation with 5% trifluoroacetic acid (TFA) for 1 h, followed by addition of 2.5% TFA/50% acetonitrile (ACN) for 1 h at 37°C. The combined supernatants were dried in a SpeedVac concentrator (Thermo Fisher Scientific, #SPD120) for mass spectrometry (MS) analysis.

### Mass spectrometry

Samples were analyzed by LC-MS using UltiMate(tm) 3000 RSLCnano System (Thermo Fisher Scientific, #ULTIM3000RSLCNANO) interfaced with Q Exactive HF. Digested samples were loaded on a fused silica trap column Acclaim PepMap 100, 0.075 mm x 2 cm (Thermo Fisher Scientific, #164569). After washing for 5 min at 5 µl/min with 0.1% TFA, the trap column was brought in-line with an analytical column (Nanoease MZ peptide BEH C18, 130A, 1.7um, 0.075 mm x250 mm, Waters, # 186008800) for LC-MS/MS. Peptides were fractionated at 300 nL/min using a segmented linear gradient from 4 to 90% B (A: 0.1% formic acid, B: 0.08% formic acid, 80% ACN): 5 min, 4-10% B; 60 min, 10-40% B; 15 min, 40-55% B; 10 min, 55-90% B. Mass spectrometry data was acquired using a data-dependent acquisition procedure with a cyclic series of a full scan acquired with a resolution of 120,000 followed by MS/MS scans (30% of collision energy in the HCD cell) with resolution of 30,000 of 20 most intense ions with dynamic exclusion duration of 20 sec.

The peak list of the LC-MS/MS were generated by Thermo Proteome Discoverer (v. 2.1) into MASCOT Generic Format (MGF) and searched against TAIR protein database (containing 35,386 sequences) using an in-house version of X! tandem (SLEDGEHAMMER (2015.09.01), the gpm.org) with carbamidomethylation on cysteine as fixed modification and oxidation of methionine as variable modification. A +/-7 ppm and 20 ppm were used as tolerance for precursor and product ions respectively. Trypsin/P was selected as the digestion enzyme and one missed cleavage per peptide was allowed. The False Discovery Rate was estimated for all samples by using a reverse database (FDR<0.01). All identified spectra belong to Arabidopsis database and peptide log (E) < = −2 (E value<=0.01) was reported. Spectral counts of the proteins in each sample were used for relative quantitation.

### Yeast two hybrid analysis

Yeast two hybrid was performed as described elsewhere^53^. Yeast cultures were grown at 28°C on Yeast Peptone Dextrose (YPD) or on selective Synthetic Dextrose (SD) media. The AH109 strain was used for transformation, following protocol described in the Yeast Protocols Handbook (version no. PR973283 21; Clontech), with both Gal4-BD and Gal4-AD constructs added. Transformants were selected on SD medium lacking Trp and Leu (SD-LW). Transformed yeast were grown to OD600 and spotted onto selective SD medium additionally lacking His (SD-LWH) which permitted the identification of weak protein interactions. Yeast dilutions during spotting were: 1/1, 1/5, 1/25, 1/25.

### Expression analysis

Total RNA extraction was performed using the hot phenol method, as described elsewhere^54^. Genomic DNA contamination was removed using TURBO DNA-free kit (Invitrogen, #AM1907) following the manufacturer’s guidelines. cDNA was synthesised using SuperScript IV First-strand Synthesis System (Invitrogen, #18091050), using gene-specific primers. cDNA was diluted 10 times before qPCR. All primers are listed in the Table S2. cDNA was amplified using SYBR Green I Master (Roche, 04887352001) and run on LightCycler 480 machine (Roche, 05015243001). Ct values were normalized to geometric mean of reference genes: *UBIQUITIN CARRIER PROTEIN 1 (UBC)* and/or *SERINE/THREONINE PROTEIN PHOSPHATASE 2A (PP2A)*. Chromatin-associated RNA has been isolated using urea as published before^55^.

### Nucleosome stability analysis

The protocol has been adapted from^56^. 2g Arabidopsis seedlings were ground in liquid nitrogen, resuspended in 35 ml NIB buffer (10 mM MES-KOH pH=5.4, 10 mM NaCl, 10 mM KCl, 2.5 mM EDTA, 250 mM Sucrose, 0.1 mM Spermine, 0.5 mM Spermidine, 1mM DTT, 1x Protease inhibitor cocktail EDTA-free (Thermo Fisher Scientific, # A32965)) and filtered through one layer of miracloth. 20% Triton X-100 was added to final concentration of 0.35%, the extract was rotated at 4°C for 10 min and spun at 1000 g for 10 min. The pellet was transferred to Low Protein Binding tubes (Thermo Fisher Scientific, #90410) and washed twice with MNase buffer (10 mM Tris-HCl pH=8, 5 mM NaCl, 2.5 mM CaCl2, 2 mM MgCl2, 1x Protease inhibitor cocktail EDTA-free (Thermo Fisher Scientific, # A32965))) at 1000g for 3 min. The pellet was weighted and MNase buffer was added in ratio 5 μL buffer:1 μg pellet. The mixture was pre-warmed at 37°C for 5 min. 150 μL of the mixture was put aside as an uncut sample and other 150 μL was subjected to 5U/ml (final concentration) MNase (Takara Bio, #2910A) enzymatic digestion at 37°C for 10 min. The concentration of MNase was experimentally tested to obtain nucleosome ladder as in^47^. The reaction was stopped by adding EGTA to final concentration of 2 mM. The mixture was spun at 1000 g for 3 min, supernatant was removed, the pellet resuspened in 80 mM salt buffer (80 mM NaCl) and rotated at 4°C for 15 min. The samples were spun at 1000 g for 3 min and the supernatant was collected as mobile nucleosome fraction. Samples were adjusted to 500 μL and subjected to DNA recovery by adding 1/50 volume 0.5 M EDTA, 1/50 vol. 5M NaCl and 2 μL RNAse A (Thermo Fisher Scientific, #EN0531). The samples were incubated at 37°C for 30 min. Proteins were removed by adding 1/16 vol. 10% SDS to final concentration 0.63% and 2.5 μL Proteinase K (Ambion, #AM2546), followed by incubation at 50°C for 1 h. The samples were then subjected to standard phenol-chloroform-isoamyl alcohol and DNA precipitation with ethanol. The enrichment was calculated by qPCR with primers indicated in Table S2. Ct values in mobile nucleosome fraction were normalized to uncut DNA and Gypsy-like transposon (At4g07700) as an internal control, following normalizations for published in^57^.

### Micrococcal nuclease assay (MNase-qPCR)

2g Arabidopsis seedlings were ground in liquid nitrogen and the pellet was resuspended in Honda buffer^21^, filtered through miracloth and spun at 1000 g, 4°C for 10 min, similarly to ChIP protocol. Subsequently, a pellet was washed twice in MNase buffer (10 mM Tris-HCl pH=8, 5 mM NaCl, 2.5 mM CaCl2, 2 mM MgCl2, 1x Protease inhibitor cocktail EDTA-free (Thermo Fisher Scientific, # A32965))) and finally resuspended in 1 mL MNase buffer. The samples were divided into 300 μL uncut control and 300 μL MNase-digested sample. The mixtures were prewarmed at 37°C for 5 min and digested samples were treated with 66.6 U/mL (final conc.) MNase enzyme (Takara Bio, #2910A) at 37°C for 30 min. The reactions were stopped by adding 1/100 volume 0.5 M EDTA and subjected to protein extraction by adding 0.05% (final conc.) SDS, followed by standard phenol-chloroform-isoamyl alcohol and DNA precipitation with isopropanol. The enrichment was calculated by qPCR with primers indicated in Table S2. Ct values in mobile nucleosome fraction were normalized to uncut DNA.

### Formaldehyde-assisted isolation of regulatory elements (FAIRE)

FAIRE was performed as shown in^57^. The enrichment was assessed by qPCR using primers indicated in Table S2. Calculations were performed using ΔΔCt method as described in^57^ with normalization of crosslinked sample (FAIRE) to non-crosslinked sample (UN-FAIRE) and then to UBC9 (AT4G27960) as an internal control.

### Chromatin immunoprecipitation

Histone modification and PolII ChIP was performed as previously described^17,55^, respectively. Immunoprecipitation was done with antibodies: anti-H3 (Abcam, #ab1791), anti-H3K27me3 (Millipore, #07-449), H2AK119ub (Cell Signalling Technology, #8240), anti-H3K36me3 (Abcam, #ab9050). PolII antibodies were as in^58^. All ChIP experiments were followed by qPCR with indicated primer pairs (Table S2).

### Statistical analysis

Statistical analyses were performed in Microsoft Excel and R Studio (R v4.0.2). P-value, p-value levels and sample number are included in figures and figure legends. Significance calculations with Student T-test were precluded by F-test to determine the usage of heteroscedastic or homoscedastic test. Unless otherwise stated, Student T-tests were two-tail. P-value levels were marked by asterisks on the figures as follows: * (p≤0.05), ** (p≤0.01), *** (p≤0.001).

## Supporting information

Mikulski et al - supplementary figures and tables

## Acknowledgements

This work was funded by the BBSRC grant BB/P006590/1, Wellcome Trust grant 210654/Z/18/Z and supported by the BBSRC Institute Strategic Programmes GRO (BB/J004588/1) and GEN (BB/P013511/1) and a Royal Society Professorship to CD. The authors would like to thank all members of the Dean lab for discussions.

## Author information

### Contributions

PM, PW and CD conceived the study. PM performed: yeast-two-hybrid, FAIRE, nucleosome mobility, ChIPs and expression analyses in *val1* and PRC1 mutants. PW performed: expression analyses, histone marks and PolII ChIPs for ASAP mutants. DZ performed nucleosome position experiment. TL performed protein co-immunoprecipitation with mass spectrometry. PM, PW and CD designed the experiments and analyzed the data. CD lead the general concept of research. PW wrote the paper together with CD with all authors contributing to different relevant sections of the manuscript.

## Ethics declarations

### Competing interest

The authors declare no competing interest.

## Additional information

Supplementary Information is available for this paper. Correspondence and requests for materials should be addressed to CD. Reprints and permissions information is available at www.nature.com/reprints.

## Notes

### Competing Interest Statement

The authors have declared no competing interest.

### Summary of Updates

Figure 1 legend has been amended to reflect the correct order of the panels.

