## Supplementary material for "VAL1 as an assembly platform co-ordinating co-transcriptional repression and chromatin regulation at Arabidopsis *FLC*": Mikulski et al - supplementary figures and tables

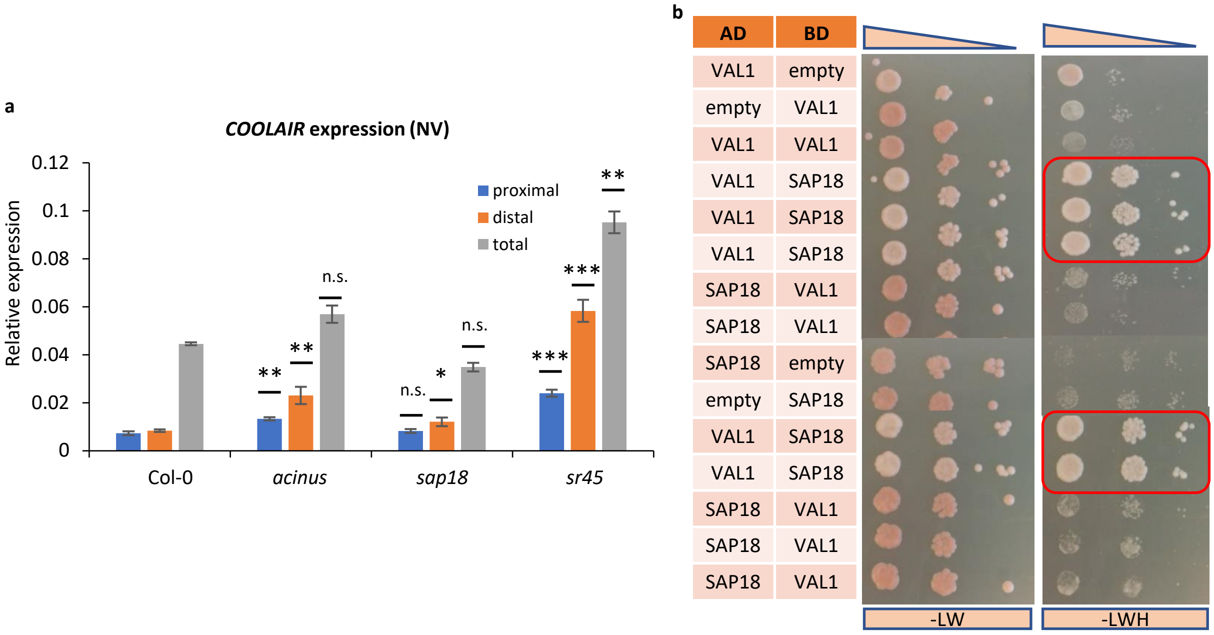


**Fig.S1. Regulation of *COOLAIR* expression by ASAP and VAL1-SAP18 interaction. a.** *COOLAIR* expression in ASAP single mutants. Y-axis corresponds to mean expression relative to *UBC*. N = 3 biological replicates; error bars = SEM; NV = non-vernalized. Statistics are in comparison to wildtype, calculated through two-tail Student T-test. **b.** . VAL1-SAP18 yeast two hybrid full results. Yeast growth was scored on selective (-LWH) medium, non-selective (-LW) medium was used as a control. Top panel shows decreasing concentration of yeast culture used for spotting. Concentration dilutions are: 1/1, 1/5, 1/25 from initial culture at OD600 = 0.8. Red frame depicts protein pairs showing yeast growth over negative controls (pairs with empty vector).

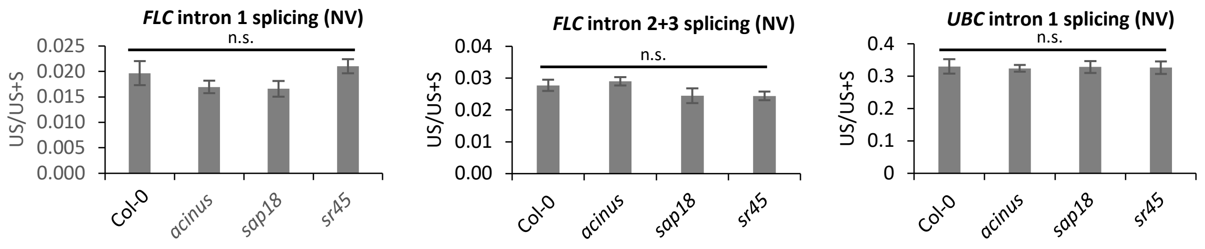


**Fig.S2. Splicing analysis in ASAP mutants**. Y-axis represents ratio unspliced (US) /total (unspliced+spliced; US+S) transcript. *FLC* intron 1 and *FLC* intron 2+3 are shown, together with *UBC* intron 1 splicing ratio used as a control. Ratio was calculated from mean expression levels normalized to *UBC*. Statistics are in comparison to wildtype, calculated through two-tail Student T-test. N = 2 biological replicates; error bars = propagated SEM; NV = non-vernalized.

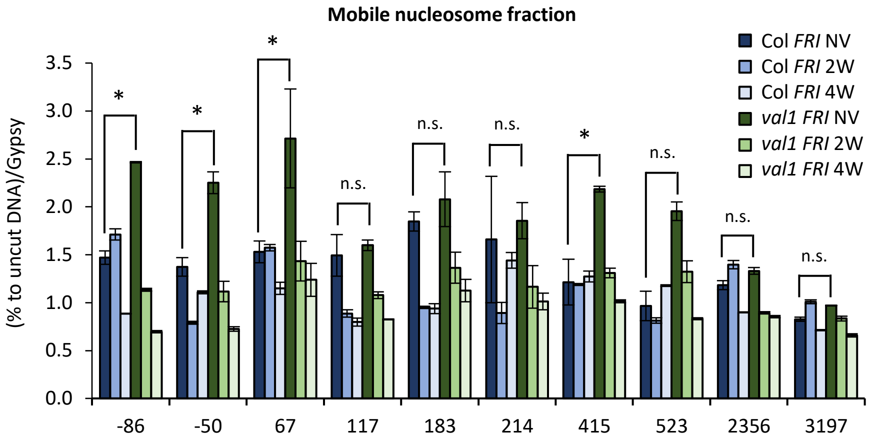


**Fig.S3. Low salt extractability of nucleosomes at *FLC*.** Shown as % recovery to uncut DNA and fold to AT4G07700 (Gypsy-like transposon). X-axis labels indicate beginning of amplicon relative to TSS. Statistics were calculated with two-tail Student T-test, in comparison *val1* to wildtype. N = 3 biological replicates; error bars = SEM; NV = non-vernalized (pre-cold); numeral-W = number of weeks of cold c VAL1-dependent change of histone marks' balance. X-axis corresponds to midpoint amplicon at *FLC*. N = 3 biological replicates; error bars = SEM; NV = non-vernalized (pre-cold); numeral-W = number of weeks of cold; T-numeral = number of days post-cold.

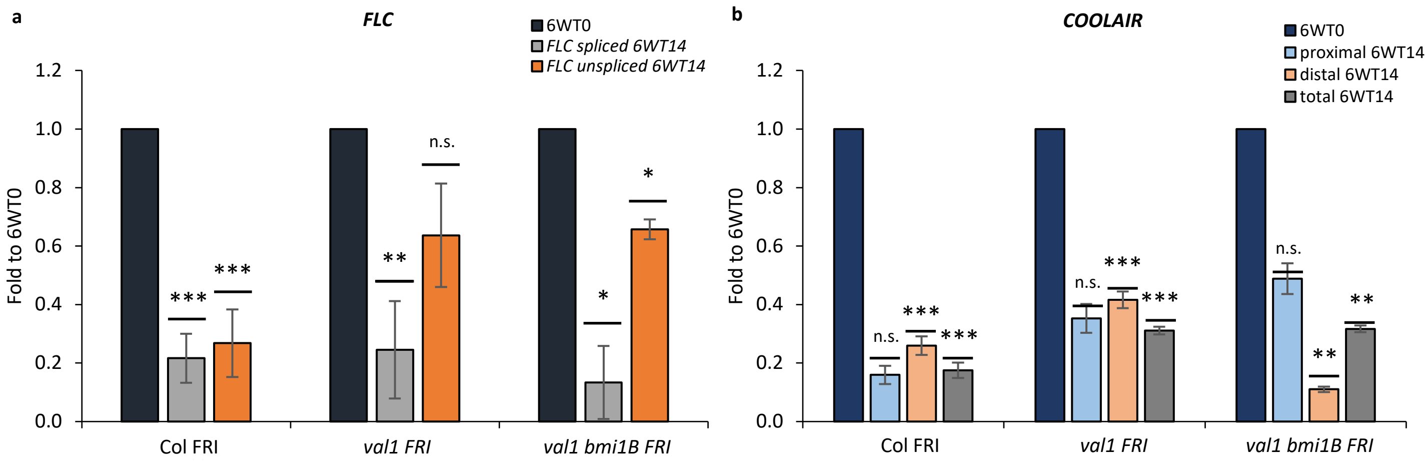


**Fig.S4. Degree of re-activation of *FLC*/*COOLAIR* expression following return to warm conditions.** Y-axis corresponds to mean expression relative to geometric mean of *UBC* and *PP2A*, normalized to expression in cold (6WT0). N = 3 biological replicates; error bars = propagated SEM; nominal-W = number of weeks of cold; T-nominal = number of days post-cold. Statistics correspond to comparison to 6WT0 timepoint of respective target RNA, calculated with two-tail Student T-test. **a.** *FLC* expression **b.** *COOLAIR* expression.

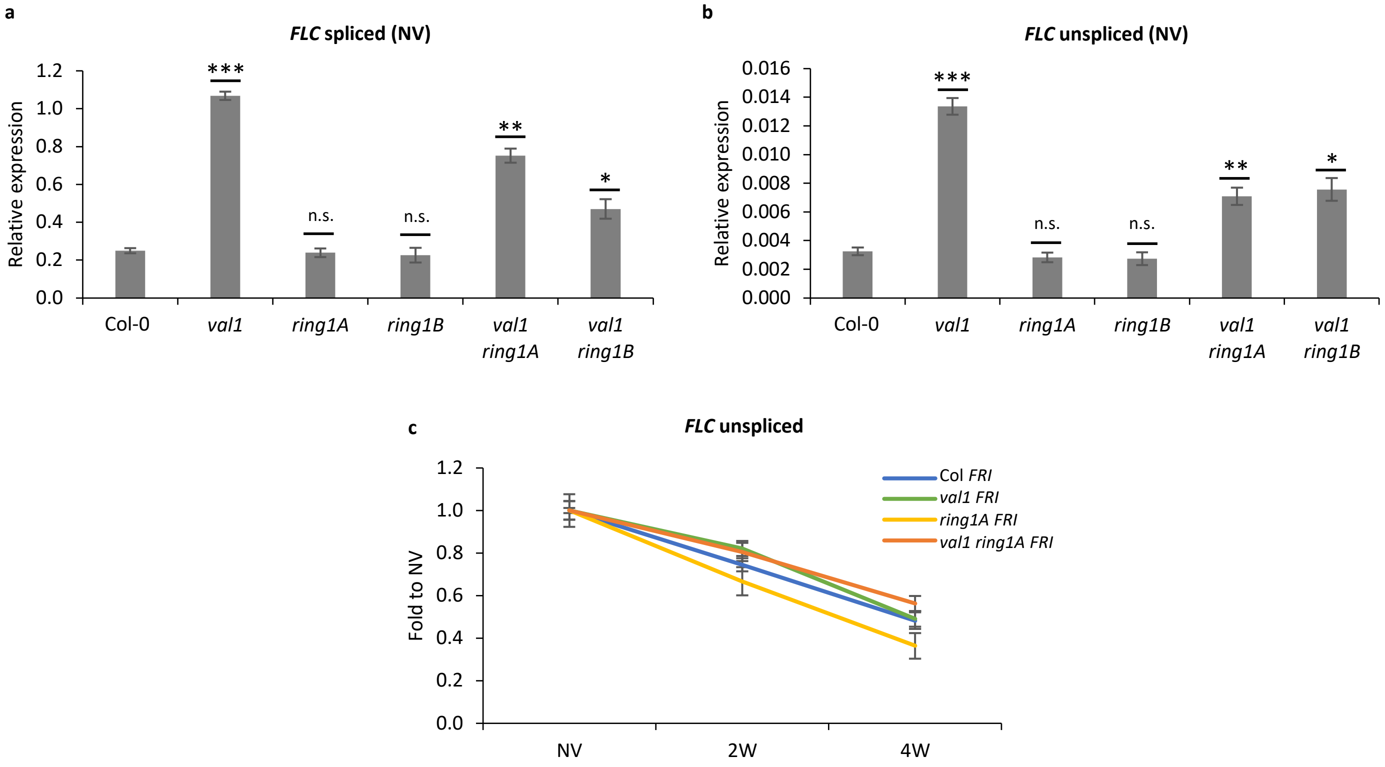


**Fig.S5. *FLC* expression in VAL1-PRC1 mutants.** Y-axis corresponds to mean relative expression to geometric mean of *UBC* and *PP2A*. N = 3 biological replicates. **a.** *FLC* spliced transcript expression **b.** *FLC* unspliced transcript expression. Statistics were calculated with Student T-test, in comparison to wildtype. **c.** *FLC* downregulation slope during vernalization (fold change to NV). Y-axis corresponds to mean relative expression to geometric mean of *UBC* and *PP2A* normalized to NV levels. N = 3 biological replicates, error bars = propagated SEM.

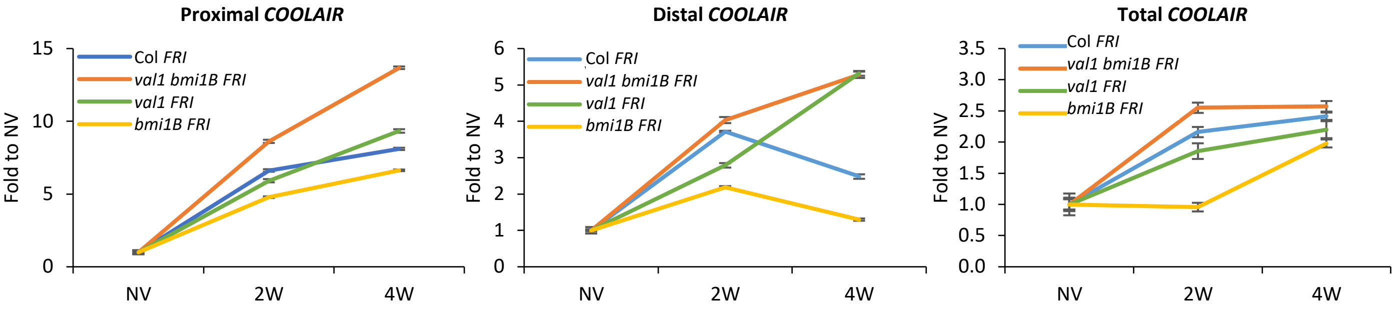


**Fig.S6. Regulation of *COOLAIR* expression by VAL1, PRC1 and NDX.** *COOLAIR* dynamics during vernalization timecourse in PRC1-VAL1 mutants. Y-axis corresponds to mean expression relative to geometric mean of *UBC* and *PP2A*, further normalized to NV levels. N = 3 biological replicates; NV = non-vernalized; nominal-W = number of weeks of cold; error bars = propagated SEM.

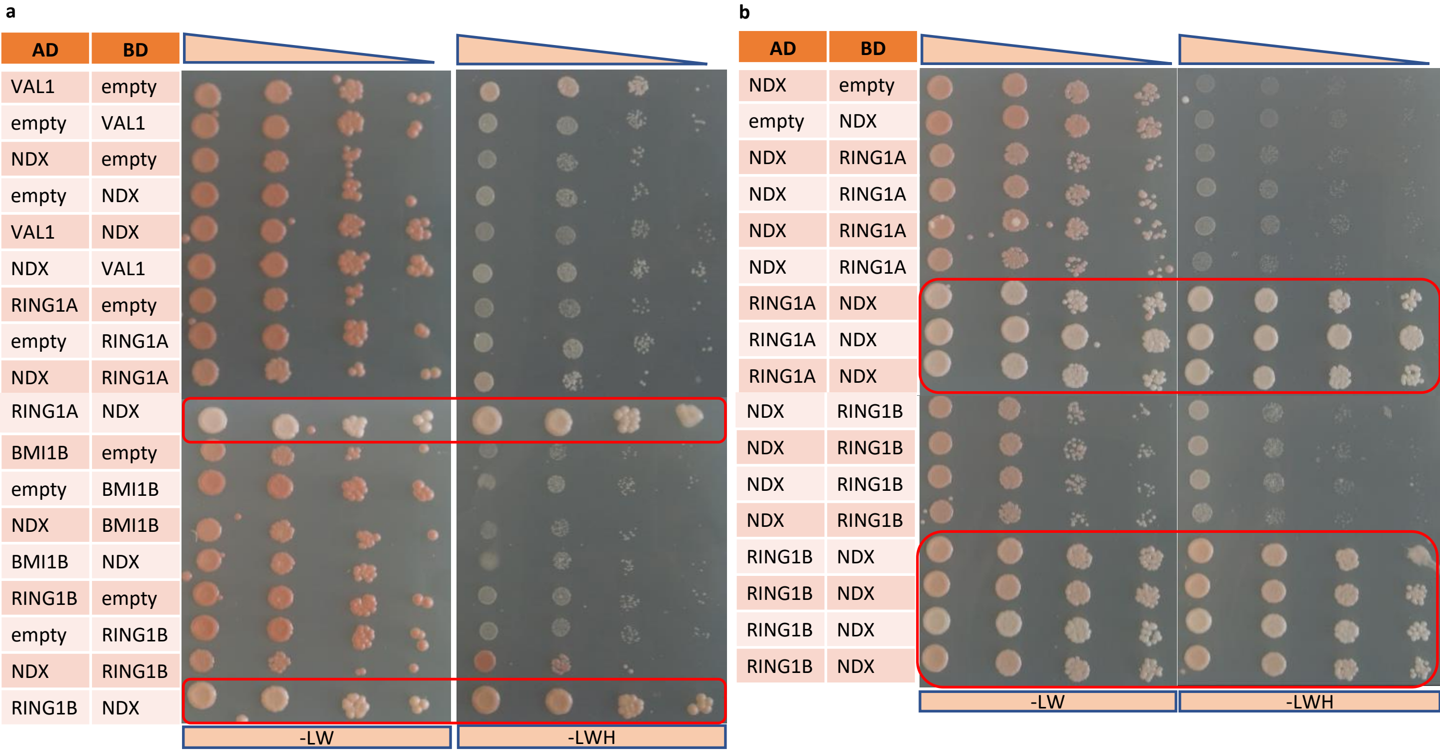


**Fig.S7. NDX-PRC1 interaction. a.** Full results on yeast two hybrid NDX-PRC1 interaction. **b.** Validation of positive interactions from a. in multiple yeast colony replicates. Yeast growth on selective medium (-LWH) and non-selective medium (-LW) is shown. Top panel corresponds to decreasing yeast culture concentration in dilutions: 1/1, 1/5, 1/25, 1/125 from initial culture at OD600 = 0.8. AD = pGAD vector backbone with Gal4 activating domain; BD = pGBKT vector backbone with Gal4 binding domain; "empty" = empty vector without insertion as negative control. Red frame depicts protein pairs showing yeast growth over negative controls (pairs with empty vector).

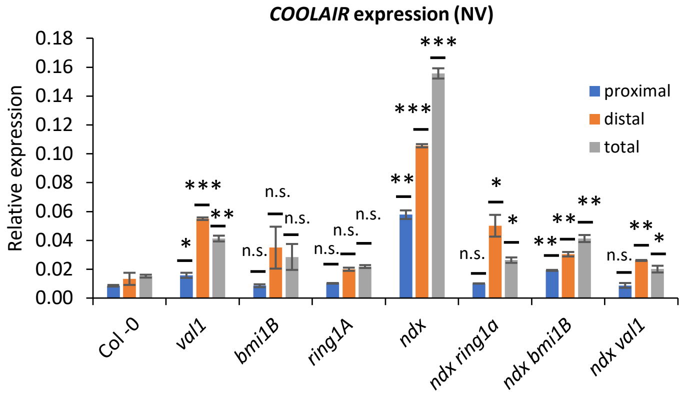


**Fig.S8. Regulation of *COOLAIR* expression by VAL1, PRC1 and NDX.** COOLAIR expression in PRC1 single mutants and NDX-PRC1 double mutants. Y-axis corresponds to mean expression relative to geometric mean of *UBC* and *PP2A*. Statistics were calculated through two-tail Student T-test. N = 3 biological replicates; error bars = SEM; NV = non-vernalized.

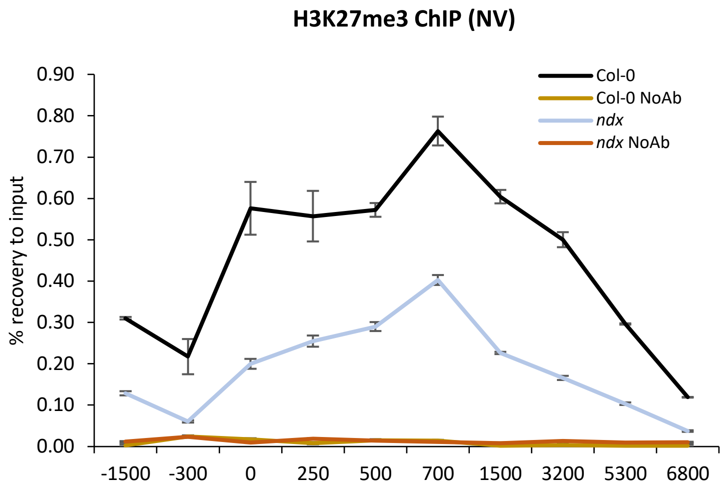


**Fig.S9. H3K27me3 enrichment in *ndx*.** H3K27me3 ChIP results in wildtype and ndx. X-axis represents midpoint of the amplicons over *FLC*. NoAb = no antibody negative control; N = 2 biological replicates for H3K27me3 IPs and NoAb controls; error bars = SEM.

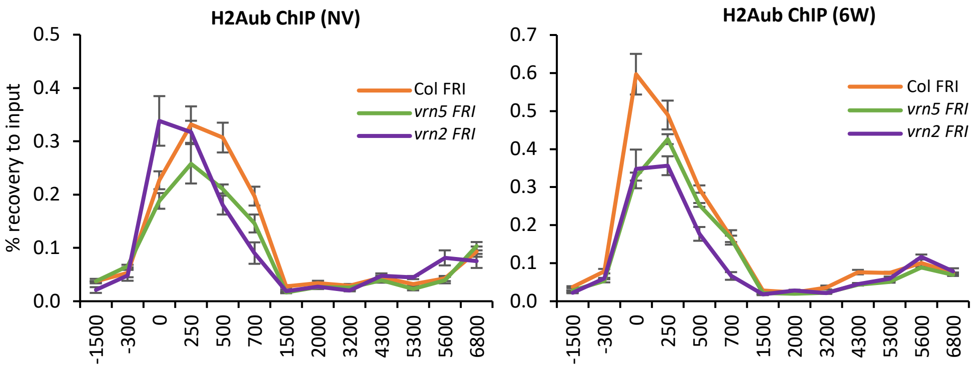


**Fig.S10. H2Aub enrichment in mutants of PRC2- and PRC2-associated components.** X-axis represents midpoint of the amplicons over *FLC*. NoAb = no antibody negative control; N = 3 biological replicates for H2Aub IPs and 3 qPCR technical replicates for NoAb controls; error bars = SEM.

**Table S1. Top enriched proteins from VAL1::VAL1-HA IP-MS/MS**

| Locus | Uniprot ID | Name | Abbreviation | IP1 (-) | IP2 (-) | IP1 (-) | IP2 (-) | IP1 | IP1 | IP2 | IP1 | IP2 |
|---|---|---|---|---|---|---|---|---|---|---|---|---|
| | | | | NV | NV | 4W T0 | 4W T0 | NV | 4W T0 | 4W T0 | 4W T7 | 4W T7 |
| AT2G30470 | Q8W4L5 | B3 domain transcription repressor | VAL1 | 0 | 0 | 0 | 0 | 53 | 69 | 80 | 61 | 66 |
| AT1G06770 | Q9M9Y4 | RING-type E3 ubiquitin transferase | BMI1B | 0 | 0 | 0 | 0 | 3 | 7 | 13 | 4 | 6 |
| AT2G30580 | Q94AY3 | RING-type E3 ubiquitin transferase | BMI1A | 0 | 0 | 0 | 0 | 10 | 14 | 12 | 11 | 7 |
| AT5G44280 | Q9FKW0 | RING-type E3 ubiquitin transferase | RING1A | 0 | 0 | 0 | 0 | 6 | 6 | 5 | 4 | 3 |
| AT4G03090 | F4JI44 | Nodulin homeobox | NDX | 0 | 0 | 0 | 0 | 3 | 8 | 13 | 7 | 9 |
| AT4G20890 | P29517 | Tubulin beta-9 | TUB9 | 0 | 0 | 0 | 0 | 21 | 11 | 0 | 18 | 14 |
| AT5G39850 | Q9FLF0 | 40S ribosomal protein S9-2 | | 0 | 0 | 0 | 0 | 11 | 8 | 0 | 6 | 0 |
| AT5G20000 | Q94BQ2 | 26S proteasome AAA-ATPase subunit | RPT6b | 0 | 0 | 0 | 0 | 7 | 8 | 0 | 10 | 0 |
| AT1G15250 | Q8LFH7 | eL20 ribosomal protein | RPL37A | 0 | 0 | 0 | 1 | 6 | 5 | 1 | 5 | 1 |
| AT2G21130 | Q9SKQ0 | Cyclophilin-like peptidyl-prolyl cis-trans isomerase | CYP19 | 0 | 0 | 0 | 0 | 5 | 7 | 0 | 7 | 0 |
| AT5G18190 | Q9FK52 | MUT9-like casein kinase | PPK2 | 0 | 0 | 0 | 0 | 5 | 4 | 0 | 4 | 0 |
| AT4G35800 | P18616 | DNA-directed RNA | NRPB1 | 0 | 0 | 0 | 0 | 4 | 5 | 3 | 6 | 1 |

| | | | | | | | | | | | | |
|---|---|---|---|---|---|---|---|---|---|---|---|---|
| | | polymerase II subunit 1 | | | | | | | | | | |
| AT4G10450 | Q9SZX9 | 60S ribosomal protein L9 | | 0 | 0 | 0 | 0 | 11 | 12 | 16 | 10 | 0 |

**Table S2. Oligonucleotides used in the study. Number in amplicon name indicates approx. midpoint position of amplicon to FLC TSS**

| Amplicon name | Forward (5'->3') | Reverse (5'->3') |
|---|---|---|
| **Primers for ChIP** | | |
| FLC -2300 | ATCCAGAAAAGGGCAAGGAG | CGAATCGATTGGGTGAATG |
| FLC -1500 | TGGAGGGAACAACCTAATGC | TCATTGGACCAAACCAAACC |
| FLC -300 | ACTATGTAGGCACGACTTTGGTAAC | TGCAGAAAGAACCTCCACTCTAC |
| FLC 0 | GCCCGACGAAGAAAAAGTAG | TTCAAGTCGCCGGAGATACT |
| FLC 250 | CTGTTCTCTGTGACGCATCC | AGGGGGAACAAATGAAAACC |
| FLC 500 | GGCGGATCTCTTGTTGTTTC | CTTCTTCACGACATTGTTCTTCC |
| FLC 700 | TGAAGTTTCAAGCCATCTTTGA | TCACTCTGAAAAGAGACATTAATCA |
| FLC 750 | CGTGCTCGATGTTGTTGAGT | TCCCGTAAGTGCATTGCATA |
| FLC 1200 | CCTTTTGCTGTACATAAACTGGTC | CCAAACTTCTTGATCCTTTTTACC |
| FLC 1500 | TTGACAATCCACAACCTCAATC | TCAATTTCCTAGAGGCACCAA |
| FLC 2000 | AGCCTTTTAGAACGTGGAACC | TCTTCCATAGAAGGAAGCGACT |
| FLC 2500 | AGTTTGGCTTCCTCATACTTATGG | CAATGAACCTTGAGGACAAGG |
| FLC 3200 | GGGGCTGCGTTTACATTTTA | GTGATAGCGCTGGCTTTGAT |
| FLC 4300 | AGAACAACCGTGCTGCTTTT | TGTGTGCAAGCTCGTTAAGC |
| FLC 5200 | CCGGTTGTTGGACATAACTAGG | CCAAACCCAGACTTAACCAGAC |
| FLC 5300 | TTTTTGTTATGGTTAGGTTTGGA | AGTAGCACTACTTCTAGACACTTGGA |
| FLC 5500 | AGATTATAGATACTGCTTCCAAACT | TTCACACCACCAAATAACAAC |
| FLC 5600 | TAATCATCATGTGGGAGCAG | GGAGAGTCACCGGAAGATTG |
| FLC 6000 | CGTGTGAGAATTGCATCGAG | AAAAACGCGCAGAGAGAGAG |
| FLC 6800 | TTGTAAAGTCCGATGGAGACG | ACTCGGCGAGAAAGTTTGTG |
| STM | GCCCATCATGACATCACATC | GGGAACTACTTTGTTGGTGGTG |
| ACT | GATATTCAGCCACTTGTCTGTG | CTTACACATGTACAACAAAGAAGG |
| **Primers for DRIPc** | | |
| FLC 3643 | TGAAATGTTACGAATACTAGCGTGT | GGATCAAAACTACTAGCTAACCCTTG |
| FLC 5030 | CCGGTTGTTGGACATAACTAGG | CCAAACCCAGACTTAACCAGAC |
| FLC 5327 | TTTTTGTTATGGTTAGGTTTGGA | AGTAGCACTACTTCTAGACACTTGGA |
| FLC 5442 | AGATTATAGATACTGCTTCCAAACT | TTCACACCACCAAATAACAAC |
| FLC 5531 | TGGTTGTTATTTGGTGGTGTG | ATCTCCATCTCAGCTTCTGCTC |
| FLC 5672 | CCTGCTGGACAAATCTCCGA | GGATTTTGATTTCAACCGCCGA |
| FLC 5801 | TTATTCCGCTGATAAGGGCGAG | AAGGTACAAAGTTCATCAACC |
| FLC 5948 | CGTGTGAGAATTGCATCGAG | AAAAACGCGCAGAGAGAGAG |
| FLC 6066 | CGTGTGAGAATTGCATCGAG | AAAAACGCGCAGAGAGAGAG |
| **Primers for Histone salt fractionation and MNase assay** | | |
| FLC -86 | CACTCTCGTTTACCCCCAAA | TCCTTTTCTCGCTTTATTTCTTTC |
| FLC -50 | GCCCGACGAAGAAAAAGTAG | TTCAAGTCGCCGGAGATACT |
| FLC 67 | AGGATCAAATTAGGGCACAAA | TCAATTCGCTTGATTTCTAGTTTTTT |
| FLC 117 | AAAAAACTAGAAATCAAGCGAATTG A | CTTTCTCGATGAGACCGTT |
| FLC 183 | AACGGTCTCATCGAGAAAG | GGAGAAGCTGTAGAGCTTGC |
| FLC 214 | CTGTTCTCTGTGACGCATCC | AGGGGGAACAAATGAAAACC |
| FLC 415 | GGCGGATCTCTTGTTGTTTC | CTTCTTCACGACATTGTTCTTCC |
| FLC 523 | GCTTTTTAGCTTCTTACTTTTGTTCA | TCGTGAATGACATGCAATTTT |

| | | |
|---|---|---|
| FLC 2356 | AGTTTGGCTTCCTCATACTTATGG | CAATGAACCTTGAGGACAAGG |
| FLC 3197 | GGGGCTGCGTTTACATTTTA | GTGATAGCGCTGGCTTTGAT |
| AT4G07700 (Gypsy) | CGGCCAAACTCAATGTAAGC | TCCCTCTTCTAGAGGTTTTGTCC |
| **Primers for Chromatin-associated RNA expression** | | |
| FLC 130 | ATTAGGGCACAAAGCCCTCT | CGACGTTTGGAGAAGGTGAC |
| FLC 151 | TGAGGATCAAATTAGGGCACA | GGATGCGTCACAGAGAACAG |
| FLC 256 | TCATCGAGAAAGCTCGTCAG | GAAAACCCAGGTAAGGAAAAGG |
| FLC 274 | CTGTTCTCTGTGACGCATCC | AGGGGGAACAAATGAAAACC |
| FLC 290 | GTCGCTCTTCTCGTCGTCTC | CAGAAGATAAAAGGGGGAACAA |
| FLC 371 | TTTTCATTTGTTCCCCCTTT | AGAGATCCGCCGGAAAAA |
| FLC 470 | GGCGGATCTCTTGTTGTTTC | CTTCTTCACGACATTGTTCTTCC |
| FLC 726 | TGAAGTTTCAAGCCATCTTTGA | TCACTCTGAAAAGAGACATTAATCA |
| FLC 894 | TGCTATGGGGTTAATGCTGA | GGTCCACAGCAAAGATAGGAA |
| FLC 1050 | TTTCATACACAGTAGTTTTGAATTTTG | GAATCGCAATCGATAACCAGA |
| FLC 1207 | TTGCTGTACATAAACTGGTCTAATTTT | TCCTTTTTACCATTAACCTCATACTAA |
| FLC 1393 | AACGAATTTCTCTCCTTTTTATGG | TGTAACTGCAAGAGTGGGAAA |
| FLC 1898 | AGTAGTTTGGCCATGTTGGT | TCAGGTGTCTCGACAATTCC |
| FLC 2522 | AGTTTGGCTTCCTCATACTTATGG | CAATGAACCTTGAGGACAAGG |
| FLC 3257 | GGGGCTGCGTTTACATTTTA | GTGATAGCGCTGGCTTTGAT |
| FLC 3657 | AAAAGTGGAAATTCAGATGTGCT | TTGAAAAGGCCACTGGAAAC |
| FLC 5155 (spliced transcript) | AGCCAAGAAGACCGAACTCA | TTTGTCCAGCAGGTGACATC |
| eIF1a-intron3 | ATGGTGACGCTGGTATGGTT | TCCTTCTTGTCCACGCTCTT |
| **Primers for total RNA expression** | | |
| FLC unspliced | CGCAATTTTCATAGCCCTTG | CTTTGTAATCAAAGGTGGAGAGC |
| FLC spliced | AGCCAAGAAGACCGAACTCA | TTTGTCCAGCAGGTGACATC |
| COOLAIR total | TGCATCGAGATCTTGAGTGTATGT | ACGTCCCTGTTGCAAAATAAGC |
| COOLAIR proximal | CCTGCTGGACAAATCTCCGA | TCACACGAATAAGGTGGCTAATTAAG |
| COOLAIR distal | GTATCTCCGGCGACTTGAAC | GGATGCGTCACAGAGAACAG |
| UBC | CTGCGACTCAGGGAATCTTCTAA | TTGTGCCATTGAATTGAACCC |
| PP2A | ACTGCATCTAAAGACAGAGTTCC | CCAAGCATGGCCGTATCATGT |
| **Primers for splicing ratio** | | |
| FLC intron1 unspliced | TTCTCCAAACGTCGCAACGGTCTC | CTCAGAAAAGTAAAAGAGCACAAAACAG |
| FLC intron1 spliced | TTCTCCAAACGTCGCAACGGTCTC | CATGCTGTTTCCCATATCGATCAAG |
| UBC9 intron1 unspliced | TTTGGATCTTCTTCCCGTCTT | AATCCCACGATCCAAATTCC |
| UBC9 intron1 spliced | CGTGAATTCGGAAGTCTTCAA | GCGCTACATGAAGTAGGAGGA |
| FLC intron 2/3 unspliced | CGCAATTTTCATAGCCCTTG | CTTTGTAATCAAAGGTGGAGAGC |
| FLC intron 2/3 spliced | AGCCAAGAAGACCGAACTCA | TTTGTCCAGCAGGTGACATC |
| **Primers for genotyping** | | |
| val1-2 WT | TCACAGAAGGAGCGTATTGAGT | ACCAAGACCAAGGAAGCATC |
| Salk LBb1.3 | | ATTTTGCCGATTTCGGAAC |

| | | |
|---|---|---|
| sr45-1 WT | TTTTGTTTTCCTTGTGTTGGC | GATTGGAGATCTTCTGGGAGG |
| Salk Lbb1.3 | | |
| sap18 WT | CTCAGCTACTTCTCCGACGTTAAG | AGTGAAGATTATGCTGTGAGAGGC |
| GabiKat 08474 | | |
| acinus WT | CCCAAGAACCAGCAAGATCAC | ACCCACTACAACACCAAGGT |
| Salk Lbb1.3 | | |
| bmi1B WT | ATGATGATTAAGGTGAAGAAG | CCGAGGTCGATATTGCATAC |
| Salk Lba1 | TGGTTCACGTAGTGGGCCATCG | |
| ring1A WT | AAACGTGAGAGTGTTTTGTGTTTG | AAAGCGTTTTAACAGCAACAATCT |
| GabiKat 08474 | ATAATAACGCTGCGGACATCTACAT TTT | |
| ndx1-4 WT | TTGAGGTGTGACTGATTGCC | GGCTAAGTGATAATCAGCTCTGC |
| WiscDsLox P745 | AACGTCCGCAATGTGTTATTAAGTT GTC | |
